# The *EIF4EBP1* gene encoding 4EBP1 is transcriptionally upregulated by MYC and linked to shorter survival in medulloblastoma

**DOI:** 10.1101/2024.03.06.583558

**Authors:** Laura Hruby, Katerina Scharov, Alberto Delaidelli, Daniel Picard, Christopher Dunham, Oksana Lewandowska, Tobias Reiff, Magalie Larcher, Celio Pouponnot, Poul HB Sorensen, Barak Rotblat, Guido Reifenberger, Marc Remke, Gabriel Leprivier

## Abstract

Medulloblastoma (MB) is the most common malignant brain tumor in childhood and is stratified into four molecular groups ‒ WNT, SHH, Group 3 and Group 4. Group 3 MB patients exhibit the poorest prognosis, with a 5-year overall survival of <60%, followed by Group 4 MB patients. Apart from *MYC* amplification in a subset of Group 3 MBs, the molecular pathomechanisms driving aggressiveness of these tumors remain incompletely characterized. The gene encoding the mTOR substrate and mRNA translation inhibitor eukaryotic translation initiation factor 4E-binding protein 1 (*EIF4EBP1*) represents a possible MYC target gene whose corresponding protein, 4EBP1, was shown to be more active in Group 3 versus Group 4 MBs. However, the prognostic role of 4EBP1 in MB and the mechanisms supporting 4EBP1 overexpression in Group 3 MB are still elusive. We analyzed *EIF4EBP1* mRNA expression in publicly available data sets and found an upregulation in MB as compared to non-neoblastic brain. *EIF4EBP1* mRNA expression levels were higher in Group 3 compared to Group 4 MBs. *EIF4EBP1* mRNA expression was correlated with *MYC* expression, most prominently in Group 3 MBs. Survival analyses highlighted that high *EIF4EBP1* mRNA expression was associated with reduced overall and event-free survival across all MB patients and in Group 3/Group 4 MB patients. Immunohistochemical evaluation of 4EBP1 protein expression in MB tissues confirmed that high levels of 4EBP1 are associated with poor outcome. Functional analyses revealed that MYC directly regulates *EIF4EBP1* promoter activity, providing a mechanism for increased *EIF4EBP1* mRNA levels in Group 3 MBs. Finally, we observed that 4EBP1 may support colony formation of *in vitro* cultured MB cells. Our data highlight that transcriptional upregulation of *EIF4EBP1* by MYC promotes *in vitro* tumorigenicity of MB cells and associates with shorter survival of MB patients.

## INTRODUCTION

Medulloblastoma (MB) is the most common malignant tumor of the central nervous system in children aged between 1-9 years [49]. MBs have been stratified into four distinct molecular groups, namely WNT, Sonic Hedgehog (SHH), Group 3 and Group 4, that are driven by different molecular pathomechanisms and characterized by distinct DNA methylome and gene expression profiles [61]. These four MB groups have been incorporated in the World Health Organization (WHO) Classifications of Tumors of the Central Nervous System in 2016 [39] and in 2021 [40], with group 3 and 4 MBs considered together as MBs without WNT and SHH activation (MB, non-WNT/non-SHH). Each MB group is associated with different prognosis, i.e., patients with WNT MB have the best prognosis, with a survival rate of >95% beyond 5 years, as these tumors rarely present with metastatic spread at diagnosis and respond well to current therapy. SHH MB patients present with an intermediate to poor prognosis, depending on patient age, tumor histology, and metastatic status [46]. While survival of Group 4 MB patients is considered intermediate, Group 3 MB is the most aggressive MB group, characterized by a high incidence of cerebrospinal fluid (CSF) metastasis at diagnosis and displaying a 5-year overall survival of <60% [46]. The standard treatment of MB consists of surgical resection followed by radiation and chemotherapy. Based on the MB group assignment and risk assessment, the radiation intensity is adapted and additional agents such as SHH inhibitors or novel agents are evaluated in clinical trials [46].

In contrast to SHH and WNT MB, Group 3 and Group 4 MB do not harbour frequent and well-defined genomic alterations [46]. Furthermore, the biological mechanisms underlying the difference in prognosis between Group 3 and Group 4 MB patients remain to be explained at the molecular level [45]. Certain high-risk factors, such as *MYC* amplification in Group 3 (17% of patients) or *MYCN* amplification in Group 4 (6% of patients) are recognized as important features [44], but the majority of patients in either group do not harbour these genetic alterations [44]. Further genomic analyses have been conducted to better delineate MB group-specific features. Specifically, Group 3 and Group 4 MBs were each subdivided in three subgroups ‒ namely alpha, beta and gamma [7]. Another study separated Group 3 and Group 4 MBs into eight molecular subtypes (I-VIII) by pairwise sample similarity analysis of DNA methylation profiles of a large MB patient cohort [44]. These analyses revealed that only certain subgroups or subtypes are characterized by *MYC* or *MYCN* amplification, which is associated with poor clinical outcome [7, 44]. In particular, only the Group 3 subgroup gamma exhibited a gain or an amplification of *MYC* [7], which was associated with the poorest overall survival among the Group 3 subgroups [7]. *MYCN* amplification was mainly detected in Group 4 subgroup alpha MBs [7]. In molecular subtypes as defined by Northcott *et al.*, *MYC* amplification is more frequent in subtypes II and III, which include Group 3 MB patients only, and in subtype V, consisting mostly of Group 4 but also include Group 3 tumors [44, 46]. Stratification of Group 3 and Group 4 MB patients has been recently harmonized by analyzing a large number of MB patients, including patients of the Cavalli *et al.* cohort [59]. This highlighted the same eight subtypes as initially defined by Northcott *et al.* [44, 59], which are now incorporated as eight subgroups (I-VIII) in the 2021 WHO classification [40].

In mice, *MYC* overexpression, together with *Trp53* deletion, drives initiation and supports maintenance of MBs that resemble human Group 3 MBs [29, 52], highlighting the contribution of MYC to MB pathogenesis and aggressiveness. Paradoxically, *MYC* mRNA expression is also elevated in the WNT MB group, to a similar level as in Group 3 MBs, which indicates that *MYC* expression – in contrast to *MYC* amplification – is not a reliable prognostic factor in MB patients [57]. As a transcription factor, MYC regulates the expression of numerous pro-tumorigenic genes [14]. One such MYC target gene, with potential clinical relevance in MB, is the eukaryotic initiation factor 4E binding protein 1 (*EIF4EBP1*) [60, 2].

*EIF4EBP1* encodes the mRNA translation inhibitor 4EBP1, which is directly regulated by the energy-sensing mechanistic target of rapamycin complex 1 (mTORC1) [43]. While under normal conditions 4EBP1 is phosphorylated and blocked by mTORC1, 4EBP1 gets activated under metabolic stress conditions following mTORC1 inhibition, and thus binds and blocks the mRNA translation initiation factor eIF4E, leading to inhibition of mRNA translation initiation [37, 63]. While 4EBP1 appears to exert tumor suppressor activity, since it blocks the oncoprotein eIF4E [33], inhibits cellular proliferation [19] and restricts tumor growth in genetically engineered mouse models of prostate [18] as well as head and neck squamous cell carcinoma (HNSCC) [65], pro-tumorigenic functions also have been reported for 4EBP1. Indeed, it was reported that 4EBP1 promotes angiogenesis in ovarian and breast cancer models, thereby facilitating tumor growth under hypoxia [6, 34], supports oncogenic transformation [36, 53], and promotes glioma and Ewing sarcoma tumorigenicity [36, 21]. However, the role of 4EBP1 in MB is currently unknown.

*EIF4EBP1* mRNA expression is upregulated in numerous tumor entities [36, 67] and high *EIF4EBP1* mRNA levels correlate with poor survival in several cancer types [67, 36, 21, 60, 64, 28, 58, 8, 9]. In MB, the amount of phosphorylated 4EBP1, i.e., inactive 4EBP1, was reported to be lower in non-SHH/non-WNT MBs when compared to SHH and WNT MBs [66]. In another study, 4EBP1 protein levels were found to be higher in Group 3 versus Group 4 MBs without any changes in phosphorylated 4EBP1 levels [20], thus suggesting that 4EBP1 is more active in Group 3 MBs. However, it is currently unknown whether *EIF4EBP1* mRNA and 4EBP1 protein expression are associated with patient outcome in MB and what the drivers of 4EBP1 overexpression in Group 3 MBs are. So far, only few transcription factors have been characterized to promote *EIF4EBP1* transcription in other tumor entities, including the androgen receptor in prostate cancer [38], ETS1 and MYBL2 in glioblastoma [24], and MYCN in neuroblastoma [64]. Additionally, MYC was shown to directly control *EIF4EBP1* transcription in colon adenocarcinoma [60] and prostate cancer cells [2], supporting that *EIF4EBP1* represents a MYC target gene.

Here, we analysed the mRNA expression of *EIF4EBP1* in MB groups and subgroups using several publically available MB expression data sets, and assessed its potential association with *MYC* mRNA expression levels and *MYC* gene amplification status. We determined the prognostic role of *EIF4EBP1* mRNA expression in MB patients and examined 4EBP1 protein expression as a prognostic biomarker in an institutional MB patient cohort. Using functional assays, we delineated the regulation of *EIF4EBP1* transcription by MYC in MB cells and characterized the contribution of 4EBP1 to clonogenic growth of MB cells *in vitro*.

## MATERIALS AND METHODS

### Data availability and bioinformatics analysis

We obtained *EIF4EBP1* mRNA expression, co-expression and survival data from publically available non-neoplastic brain tissue and MB data sets from the R^2^ Genomic Analysis Visualization Platform (R^2^ AMC; http://r2.amc.nl). An overview of the used datasets and the corresponding GSE numbers in provided in Table S1. The number of patients per MB group in each cohort is listed in Table S2. For co-expression analyses of *EIF4EBP1* mRNA expression and *MYC* or *MYCN* mRNA expression, the Cavalli *et al.* [7] and Pfister [44] cohorts were used. *EIF4EBP1* mRNA expression data in primary and metastatic MB tissues was obtained and pooled from the Delattre, Gilbertson [55] and Thompson MB cohorts or obtained from the Cavalli *et al.* cohort [7]. Overall survival analysis was conducted using the Cavalli *et al.* [7] and Pomeroy [12] cohorts. As cut-off for distinction between high versus low expression groups, the first versus last quartile was used as cut-off across each data set.

Proteomic data were kindly provided by Dr. Ernest Fraenkel (Broad Institute of MIT and Harvard, Boston, MA) [1] and were downloaded in the original instrument vendor format from the MassIVE online repository under MSV000082644.

ChIP-seq data for MYC (UCSC Accession: wgEncodeEH001867, wgEncodeEH002800, wgEncodeEH000670, wgEncodeEH003436, wgEncodeEH001807, wgEncodeEH000547, wgEncodeEH000545, wgEncodeEH002795, wgEncodeEH000596) were downloaded from ENCODE (Encyclopedia of DNA Elements at UCSC; [13, 15]) using the human genome GRCh 38/hg 38. ChIP-seq data were obtained from ENCODE [13, 15], and included data from seven cell lines. The files were combined into a single BAM file and data where then visualized using IGV version 2.9.1 (https://igv.org; [56]).

DNA methylation data were downloaded from the GEO website (https://www.ncbi.nlm.nih.gov/geo/) for normal pediatric brain (GSE90871 [10]) and MB tissues (GSE85212 [50]). CpG sites included within nucleotides −1065 to +29,848 of *EIF4EBP1*, spanning the *EIF4EBP1* promoter region, two introns and three exons (human genome GRCh 37/hg19; Chr8: 37,886,955-37,917,868; exact chromosomal positions of the CpG sites are provided in Table S3) were selected for analysis and the mean was determined for each group and CpG site. A two-tailed Fisher’s exact test was used to determine statistical differences between normal pediatric brain tissue samples and MB groups.

Analysis of MYC target genes co-expressed with *EIF4EBP1* was performed with R^2^AMC using the Cavalli *et al*. data set [7]. Genes co-expressed with *EIF4EBP1* (cut-off: r=0.45; p<0.05) were initially identified for each MB group. GSEA was performed using the Broad 2020 09 c6 oncogenic gene set collection (p-value cut-off: p<0.05). From these genes, the ones listed in the “Broad institute: MYC_UP_V1_UP” human gene dataset [5], considered as MYC target genes, were counted and the corresponding p-value was calculated using Fisher’s exact test.

### Immunohistochemical staining for 4EBP1 protein expression

Immunohistochemistry for 4EBP1 was performed on formalin-fixed and paraffin-embedded (FFPE) MB tissue microarray (TMA) sections using standard protocols. The TMA consisted of tumor samples from 63 patients; samples were obtained after written informed consent and Institutional Review Board (IRB) approval between 1986 and 2012 from the BC Children’s Hospital (BCCH, Vancouver, British Columbia, Canada) as previously described [17, 62]. Detailed information about this cohort, including methods for subgroup assignment, was previously published [62]. Two samples were excluded due to limited amounts of remaining tissue on the TMA. Sections were deparaffinized in xylene and rehydrated over a decreasing ethanol series before being incubated in Tris EDTA buffer (CC1 standard) at 95°C for 1 hour to retrieve antigenicity. Tissue sections were then incubated with the primary antibody against 4EBP1 (Abcam ab32024, 1:200) for 1 hour (Ventana Discovery platform). Tissue sections with bound primary antibody were then incubated with the appropriate secondary antibody (Jackson antibodies at 1:500 dilution), followed by Ultramap HRP and Chromomap DAB detection. Intensity scoring was determined by an experienced pathologist and scored as positive and negative for the survival analyses. As for expression analysis in primary versus recurrent tissues, intensity scoring was performed according to a four-tiered scale: 0, no staining; 1, weakly positive staining; 2, moderately positive staining; 3, strongly positive staining. Immunohistochemical expression was quantified as H-score between 0 – 300 obtained by the product of the staining intensity (0-3) and the percentage of positive cells [41].

### Statistical analyses

Unpaired t-tests were performed when comparing gene expression (unless otherwise stated). Correlation analyses were performed by calculating Pearson correlation. GraphPad Prism version 7.04 (GraphPad Software, San Diego, CA, USA) was used for these statistical analyses. For correlative analysis of 4EBP1 staining with patient outcome, differences in survival between MB groups were calculated with log-rank tests for univariate survival analysis. Statistical analyses were performed with R 3.5.1 using the packages “survival” and “survminer” for survival analyses.

### Cell culture

HEK293-T embryonic kidney cells were obtained from American Type Culture Collections (ATCC, Manassas, VA). Med8a cells were a kind gift from Prof. Pablo Landgraf (University Hospital Cologne, Cologne, Germany), and the HD-MB03 cell line was generously provided by Prof. Till Milde (DKFZ, Heidelberg, Germany). The generation of inducible control (ishScr) and stable 4EBP1 knock-down (ish4EBP1) Med8a and HD-MB03 cells has been reported elsewhere [36]. Control and MYC-overexpressing ONS76 and UW228.3 MB cells (as described in [54]) were kindly provided by Dr. Nan Qin (University Hospital Düsseldorf, Düsseldorf, Germany). HD-MB03 cells were maintained in Roswell Park Memorial Institute (RPMI 1640) medium (61870010, Thermo Fisher Scientific, Waltham, MA, USA) supplemented with 10% fetal bovine serum (FBS) (10270-106, Thermo Fisher Scientific), 1% penicillin/streptomycin (10270-106, Sigma Aldrich, St Louis, USA) and 1% non-essential amino acids (MEM NEAA 100x) (11350912, Thermo Fisher Scientific). The remaining cell lines were maintained in Dulbecco’s modified Eagle Medium (DMEM) (10569010, Thermo Fisher Scientific) supplemented with 10% FBS and 1% penicillin/streptomycin. All cell lines were cultured in a humidified incubator at 37°C with 5% CO_2_. The cell lines were confirmed to be mycoplasma-free by Venor GeM Classic (11-1050, Minerva Biolabs, Berlin, Germany) kit and validated by STR-profiling at the Genomics & Transcriptomics Laboratory (GTL), Biological and Medical Research Center (BMFZ), Heinrich Heine University (Düsseldorf, Germany).

### siRNA transfection

Cells were transfected in 6-well plates at 70% confluency with 25 nM control siRNA (D-001206-14-50, Dharmacon, Cambridge, UK) or siRNAs targeting *MYC* (D-003282-14 & D-003282-35, Dharmacon) using siLentFect transfection reagent (1703362, Biorad, Hercules, CA, USA) (see Table S4 for siRNA sequences). Briefly, a master mix containing 125 µl Opti-MEM (31985-070, Thermo Fisher Scientific) and 3 µl siLentFect was prepared and incubated for 5 min at room temperature (RT). Meanwhile, 125 µl Opti-MEM were mixed with 25 nM of siRNA for each well. The siRNA mix was mixed 1:1 with the master mix, incubated for 20 min at RT and added dropwise onto the cells. Medium was changed the day after transfection. Cells were re-transfected after 96 h. At 168 h following the first transfection, RNA and protein were harvested for further analysis.

### Plasmid construction

The pGL4.22 plasmid containing the −192 to +1372 promoter region of the human *EIF4EBP1* gene fused to Firefly Luciferase has been reported before [64]. Each of the three identified E boxes (MYC binding sites) was mutated separately to CAAGGC. Cloning was performed by GENEWIZ Germany GmbH (Leipzig, Germany).

### Luciferase reporter assays

HEK 293-T cells were seeded in 12-well plates to reach 50% confluency at the day of transfection. Cells were transfected with 125 ng of the *EIF4EBP1* promoter Firefly luciferase plasmid (wild type or mutants), 2 ng of *Renilla* luciferase expressing pRL SV40 plasmid (E2231, Promega), as internal control, and 25, 50 ng or 100 ng of MYC expressing pcDNA3.3 MYC plasmid (kindly provided by Dr. Nan Qin, Düsseldorf, Germany), completed to 500 ng total DNA with pcDNA3.1 plasmid (V79020, Thermo Fisher Scientific) using CalFectin^TM^ Cell Transfection Reagent (SL100478, SignaGen Laboratories; Frederick, MD; USA) according to the manufacturer’s guidelines. Cells were harvested at 48 h post-transfection and activity of Firefly and *Renilla* luciferases were sequentially determined using the Dual-luciferase Reporter Assay System (E1980, Promega) and a Beckman Coulter microtiter plate reader (Beckman Coulter, Krefeld, Germany). All samples were performed in triplicate and the final luciferase quantification was formulated as the ratio of Firefly luciferase to *Renilla* luciferase luminescence. The relative luminescence was calculated by normalizing each biological replicate to either the 0 ng condition or to the *EIF4EBP1* promoter control without mutations.

### Soft agar assay

Inducible control and stable 4EBP1 knock-down HD-MB03 and Med8A cells were treated with 1 µg/ml doxycyclin 72 hrs prior to seeding. Cells were plated in 6-well plates with 8,000 cells per well in DMEM supplemented with 10% FBS in a top layer of 0.25% agar added over a base layer of 0.4% agar in DMEM supplemented with 10% FBS. Cells were fed twice a week with 1 ml of either DMEM supplemented with 10% FBS, 1% penicillin/streptomycin and 1 µg/ml doxycyclin (for Med8A) or RPMI supplemented with 10% FBS, 1% penicillin/streptomycin and 1 µg/ml doxycyclin (for HD-MB03) onto the top layer. After 3 weeks at 37°C, colonies were stained with 0.01% crystal violet and 10 random fields were counted manually for each well. The percentage of colony forming cells was calculated.

### RNA extraction, cDNA synthesis and qRT-PCR

RNA was extracted using the RNeasy Plus Mini Kit (74136, QIAgen, Hilden, Germany). The extraction was performed according to the protocol provided by the manufacturer. Isolated RNA was retro-transcribed to cDNA using 1 μg of RNA per reaction with either the QuantiTect Reverse Transcription Kit (205311, QIAgen) or the High-Capacity cDNA Reverse Transcription Kit (4368813, Applied Biosystems, Waltham, MA, USA) according to manufacturer’s protocol. Real time PCR was performed in triplicates using 1 µl cDNA and 9 µl master mix consisting of 5 µl SYBR Green PCR Mix (4309155, Applied Biosystems), 3 µl H_2_O and 1 µl of forward and reverse primers (0.5 µM final concentration). PPIA, GusB and β-actin were used as housekeepers. For primer sequences see Table S5.

### Protein extraction and immunoblot analysis

Cells were lysed in RIPA buffer (150 mM NaCl, 50 mM Tris-HCl, pH 8, 1% Triton X100, 0.5% Sodium deoxycholate, and 0.1% SDS) supplemented with proteinase inhibitor cocktail (11873580001, Roche, Basel, Switzerland) and phosphatase inhibitor (04906837001, Roche). Cell lysates were centrifuged at 14,000 x g for 15 min at 4°C and supernatants were collected. Protein concentration was quantified using the Pierce^TM^ BCA Protein Assay Kit (23225, Thermo Fisher Scientific) according to manufacturer’s protocol. Twenty micrograms of total protein were loaded either on a 12% polyacrylamide-SDS gel and transferred to a 0.2 µm nitrocellulose membrane (No10600001, GE Healthcare; Chicago, IL, USA). Membranes were blocked with 5% bovine serum albumin (BSA) (8076.3, Carl Roth, Karlsruhe, Germany) TBS-Tween (20 mM Tris-HCl, pH 7.4, 150 mM NaCl, 0.1% Tween 20) and probed with primary antibodies (as detailed in Table S6) diluted 1:1,000 in 5% BSA TBS overnight at 4°C. Membranes were then incubated with either a corresponding anti-mouse (926-32210, Li-Cor, Bad Homburg, Germany) or anti-rabbit (926-32211, Li-Cor) fluorescent secondary antibody diluted 1:10,000 or a corresponding anti-rabbit IgG, HRP linked secondary antibody (#7074, cell signaling, Cambridge, UK) diluted 1:3000. The fluorescent signal was visualized with the LI-COR Odyssey® CLx system (Li-Cor) and the chemiluminescent signal was detected using enhanced chemiluminescent reagent (ECL) and the detection device LAS-3000 mini (Fujifilm, Tokyo, Japan).

### Statistical analysis of experimental data

All experiments were carried out in three biological replicates. Data are represented as mean +/- standard deviation (SD). A two-sided Student’s t-test was used to compare differences between control and experimental groups. Results were considered as being statistically significant at p < 0.05. Statistical tests were calculated with GraphPad Prism version 7.04.

## RESULTS

### *EIF4EBP1* mRNA levels are elevated in MBs

To investigate *EIF4EBP1* mRNA expression in MB tissues, we pooled and analysed publically available data from two non-neoblastic brain and seven independent and non-overlapping MB datasets. We found that *EIF4EBP1* mRNA expression was significantly upregulated in MB tissues compared to non-neoplastic brain tissues (Fig. 1A). This was not related to a hypomethylation of *EIF4EBP1* promoter region, as DNA methylation levels of 18 CpG sites within the *EIF4EBP1* promoter region (hg19; Chr8: 37,886,955-37,917,868) were not different in normal pediatric brain tissues versus MB tumor tissues (Supplementary Fig. 1A). Assessing the association of *EIF4EBP1* mRNA expression with MB risk factors revealed that *EIF4EBP1* mRNA levels were higher in relapsed versus primary MB tissues (Supplementary Fig. 1B), while *EIF4EBP1* mRNA expression was similar in metastatic versus primary MB tissues (Supplementary Fig. 1C and D).

**Figure 1.**
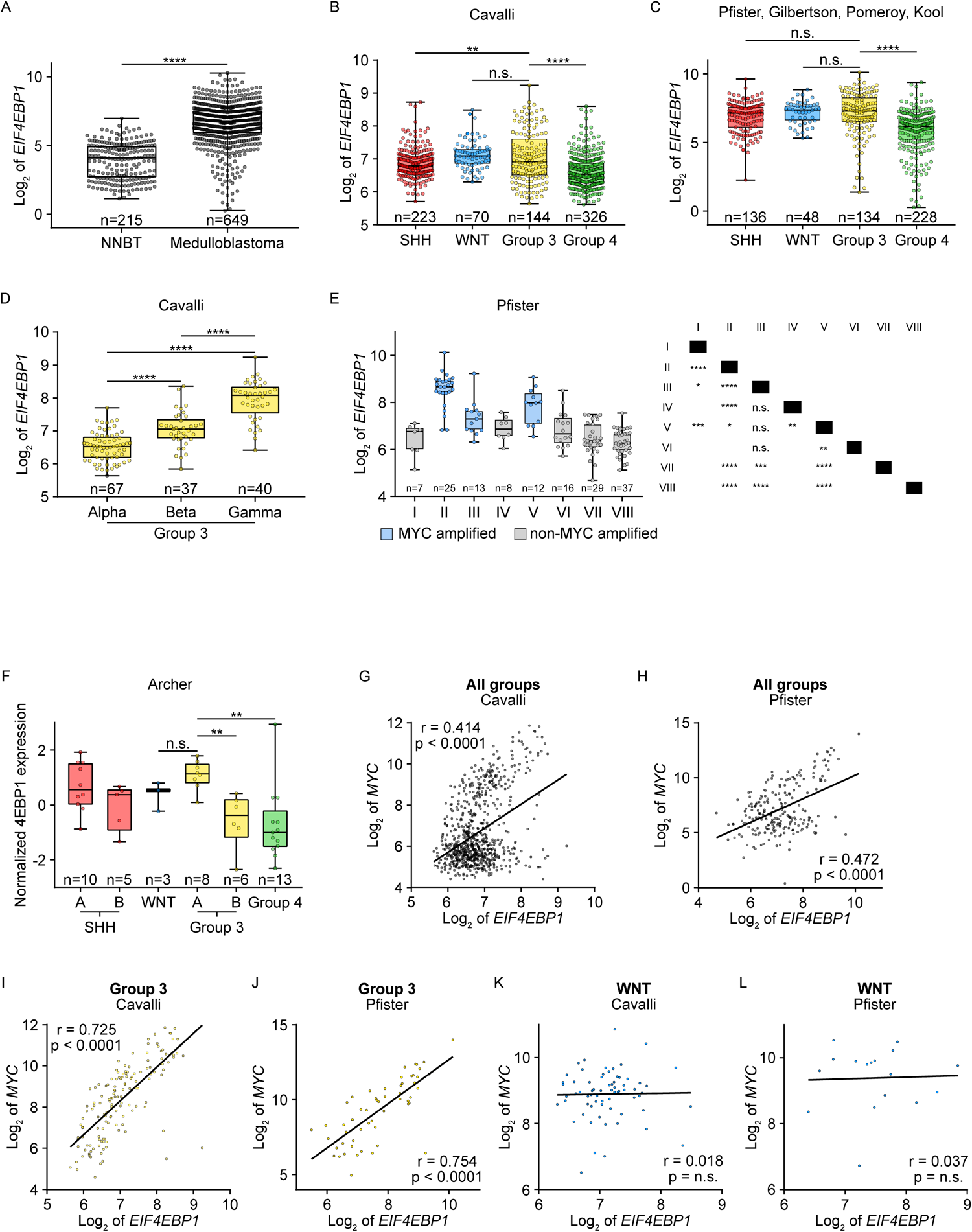
*EIF4EBP1* mRNA is upregulated and is co-expressed with *MYC* in MB. **A**, Expression levels of *EIF4EBP1* mRNA in a pool of non-neoplastic brain tissues (NNBT) (Berchtold *et al.* (n=172) [4] and Harris *et al.* (n=43) [23] cohorts) compared to a pool of MB tissues (denBoer (n=27) [16], Delattre (n=54), Gilbertson (n=73) [55], Hsieh (n=22) [25], Kool *et al.* (n=62) [30], Pfister (n=223) [44] and Pomeroy (n=188) [12] cohorts). **B and C**, Expression levels of *EIF4EBP1* mRNA according to the four MB groups SHH, WNT, Group 3 and Group 4 using the Cavalli *et al.* cohort [7] or a pool of the Kool *et al.*, Gilbertson, Pfister and Pomeroy cohorts [30, 55, 44, 12] (see Table S2 for the number of patient samples per group and Table S7 for the results of pair-wise statistic tests between different groups). **D**, Expression levels of *EIF4EBP1* mRNA according to subgroups of Group 3 MB from the Cavalli *et al.* cohort [7]. **E**, Expression levels of *EIF4EBP1* mRNA according to the Heidelberg subtypes from the Pfister cohort [44]. Significance in A-E was calculated using an unpaired and two-tailed parametric t-test (*p<0.05, **p<0.01, ***p<0.001, ****p<0.0001). **F**, Expression levels of 4EBP1 protein in MB tissues clustered to the four groups, including two subsets of SHH (A and B) and Group 3 (A and B), from [1]. p value was calculated using an unpaired and two-tailed parametric t-test (**p<0.01). **G-L**, Expression levels of *EIF4EBP1* mRNA in MB patient samples plotted against the mRNA expression levels of *MYC* in all MB patients (G and H), in Group 3 MBs (I and J) or WNT MBs (K and L) using the Cavalli *et al.* [7] and the Pfister [44] cohorts as indicated (see Table S2 for the numbers of patient samples per group). Co-expression levels were quantified by calculating the Pearson correlation coefficient.

Analysis of *EIF4EBP1* mRNA expression levels according to MB groups showed, in two single patient cohorts [7, 47] as well as in a pooled patient cohort [11, 30, 44, 55], that *EIF4EBP1* mRNA expression was elevated in Group 3 relative to Group 4 MBs (Fig. 1B and C; Supplementary Fig. 1E; Table S7), in accordance to previous observations made for 4EBP1 protein levels using proteomics data [20]. However, *EIF4EBP1* mRNA levels were as high in WNT MBs, the least aggressive MB group, as in Group 3 MBs (Fig. 1B and C; Table S7). While *EIF4EBP1* was more strongly expressed in Group 3 compared to SHH MBs in the Cavalli *et al.* MB cohort [7]] (Fig. 1B), this difference was not obvious in pooled datasets [11, 30, 44, 55]) (Fig. 1C). We next investigated levels of *EIF4EBP1* mRNA expression according to Group 3 and Group 4 subgroups as defined by Cavalli *et al.* [7] and by Northcott *et al.* (Pfister cohort) [44]. This highlighted that in Group 3, *EIF4EBP1* was more highly expressed in the gamma subgroup, the most aggressive Group 3 subgroup (corresponding mainly to subtype II [26]), as compared to alpha and beta subgroups, while in Group 4, *EIF4EBP1* levels were higher in the alpha subgroup (corresponding to subtypes V and VI [26]) compared to the other subgroups (Fig. 1D and Supplementary Fig. 1F). Noteworthy, *MYC* gain or amplification is a feature of Group 3 subgroup gamma, while *MYCN* amplification is a characteristic of Group 4 subgroup alpha [7], thus pointing to a relationship between high *EIF4EBP1* expression and *MYC(N)* amplification in MB. In line, we uncovered in the Pfister cohort [44] that *EIF4EBP1* expression is most elevated in subtypes II, III and V, which are characterized by *MYC* amplification (20% of cases for subtype II and 10% of cases for subtypes III and V [44]) (Fig. 1E). Using proteomic data extracted from the Archer *et al.* dataset [1], we confirmed that consistently with *EIF4EBP1* mRNA expression, 4EBP1 protein expression was significantly higher in Group 3A, corresponding to subtype II [44], as compared to Group 4 and Group 3B but was at the same level than in the WNT subgroup (Fig. 1F).

Finally, we assessed *EIF4EBP1* copy number alterations in Cavalli *et al.* [7] Group 3 and Group 4 MBs as a possible mechanism supporting *EIF4EBP1* overexpression. We observed a high frequency of *EIF4EBP1* gain in Group 3 subgroup gamma (corresponding mainly to subtype II [26]) (50% of cases), which was not the case in the other Group 3 subgroups (Supplementary Fig. 1G). In contrast, only around 2% of Group 4 subgroup alpha (corresponding to subtypes V and VI [26]) showed *EIF4EBP1* gain (Supplementary Fig. 1H), indicating that levels of *EIF4EBP1* expression are impacted by copy number alterations in Group 3 MB but not in Group 4 MB subgroups.

In conclusion, *EIF4EBP1* mRNA and 4EBP1 protein expression is increased in MB, in particularly in the most aggressive subgroups characterized by *MYC* or *MYCN* gene amplification.

### *EIF4EBP1* expression is associated with *MYC* expression in MB

To elucidate the possible link between *EIF4EBP1* and *MYC* mRNA expression in MB, we analysed their expression levels in the different MB groups of the Cavalli *et al.* [7] and Pfister [44] cohorts. We found that *EIF4EBP1* and *MYC* mRNA levels were strongly associated with each other across all MB patients ([r] 0.414, p value < 0.0001; Fig. 1G and [r] 0.472, p value < 0.0001; Fig. 1H). Correlative analyses according to MB groups showed in Group 3 an even stronger correlation between *EIF4EBP1* and *MYC* mRNA expression ([r] 0.725, p value < 0.0001; Fig. 1I; [r] 0.754, p value < 0.0001; Fig. 1J). In Group 4 MBs, *EIF4EBP1* mRNA expression was not correlated with *MYC* mRNA expression ([r] 0.118, p value < 0.01; Supplementary Fig. 1I; ([r] 0.194, p value = n.s.; Supplementary Fig. 1J), but strongly associated with *MYCN* mRNA expression ([r] 0.534, p value < 0.0001; Supplementary Fig. 1K; [r] 0.507, p value < 0.0001; Supplementary Fig. 1L), consistent with *MYCN* amplification being a common hallmark feature of this MB group [46]. Further analyses indicated that *MYCN* mRNA levels neither correlated positively with *EIF4EBP1* mRNA levels across all MB groups ([r] 0.220, p value < 0.0001; Supplementary Fig. 2A; [r] 0.087, p value = n.s.; Supplementary Fig. 2B) nor in Group 3 MBs ([r] −0.199, p value < 0.05; Supplementary Fig. 2C; [r] −0.468, p value < 0.001; Supplementary Fig. 2D).

There was no significant association between mRNA levels of *EIF4EBP1* and *MYC* in the WNT MB group ([r] 0.018, p value = n.s.; Fig. 1K; [r] 0.037, p value = n.s.; Fig. 1L), even though this group displayed high *MYC* mRNA expression levels in the analysed cohorts [57]. Moreover, only five MYC target genes (as defined from the human gene set “Broad institute: MYC_UP.V1_UP” [5]) were significantly co-expressed with *EIF4EBP1* in WNT MBs as opposed to 56 MYC target genes in Group 3 MBs (Table 1). These data are in accordance with the findings of Forget *et al.* [20], namely that MYC activity is lower in WNT MBs as in Group 3 MBs despite similar levels of *MYC* transcripts. Further analysis of the Cavalli *et al.* [7] and Pfister [44] cohorts highlighted highly significant associations between *EIF4EBP1* and *MYC* mRNA levels in the *MYC* amplified MB subgroups Group 3 subgroup gamma (corresponding mainly to subtype II [26]) ([r] 0.704, p value < 0.0001; Supplementary Fig. 2E) and subtypes II, III and V combined ([r] 0.604, p value < 0.0001; Supplementary Fig. 2F), confirming co-expression of *EIF4EBP1* and *MYC* in MYC-driven MB tissues.

**TABLE 1:** GSEA of MYC-upregulated genes co-expressed with *EIF4EBP1* in MB groups.

| Group | p-value | Number of genes |
| --- | --- | --- |
| SHH | 6.2e-11 | 16 |
| WNT | 0.03 | 5 |
| Group 3 | 4.8e-45 | 56 |
| Group 4 | 0.03 | 3 |

Broad institute: MYC\_UP.V1\_UP; cut-off: r=0.45

### High *EIF4EBP1* mRNA expression is associated with shorter survival of MB patients

We next determined whether *EIF4EBP1* mRNA expression is linked to prognosis of MB patients. To do so, we analysed two independent and non-overlapping MB patient cohorts, i.e., Cavalli *et al.* [7] and Pomeroy [12] cohorts. Kaplan Meier estimates revealed that high *EIF4EBP1* mRNA levels (using first versus last quartile of expression level as cut-off) was significantly associated with reduced overall survival across all MB groups in both cohorts (p value = 0.013; Fig. 2A; p value = 9.3e^-03^; Fig. 2B). When restricting our analyses to the most aggressive cases, focusing on Group 3 and Group 4 patients combined, we uncovered that high *EIF4EBP1* expression was similarly associated with poor outcome in both cohorts (p value = 2.8e^-03^; Fig. 2C; p value = 7.9e^-03^; Fig. 2D). However, in the same cohorts there was no significant association between *EIF4EBP1* mRNA levels and overall survival in Group 4 MB patients only (p value = 0.096; Supplementary Fig. 3A; p value = 0.184; Supplementary Fig. 3B). This is in contrast with the association we observed between high *EIF4EBP1* expression and unfavorable outcome in Group 3 MB patients of the Cavalli *et al.* cohort [7] (p value = 0.025; Fig. 2E). Further analyses indicate that *EIF4EBP1* mRNA expression levels were not correlated to overall survival in the WNT MB group (p value = 1.000; Fig. 2F) and the SHH MB group (p value = 0.272; Supplementary Fig. 3C) of the Cavalli *et al.* cohort [7]. However, *EIF4EBP1* expression could not be analysed using the Pomeroy *et al.* cohort [12] for the Group 3, WNT or SHH MB groups separately as patient numbers were too low. These data suggest that high *EIF4EBP1* expression are linked to less favorable prognosis across all MB patients as well as in Group 3 MB patients.

**Figure 2.**
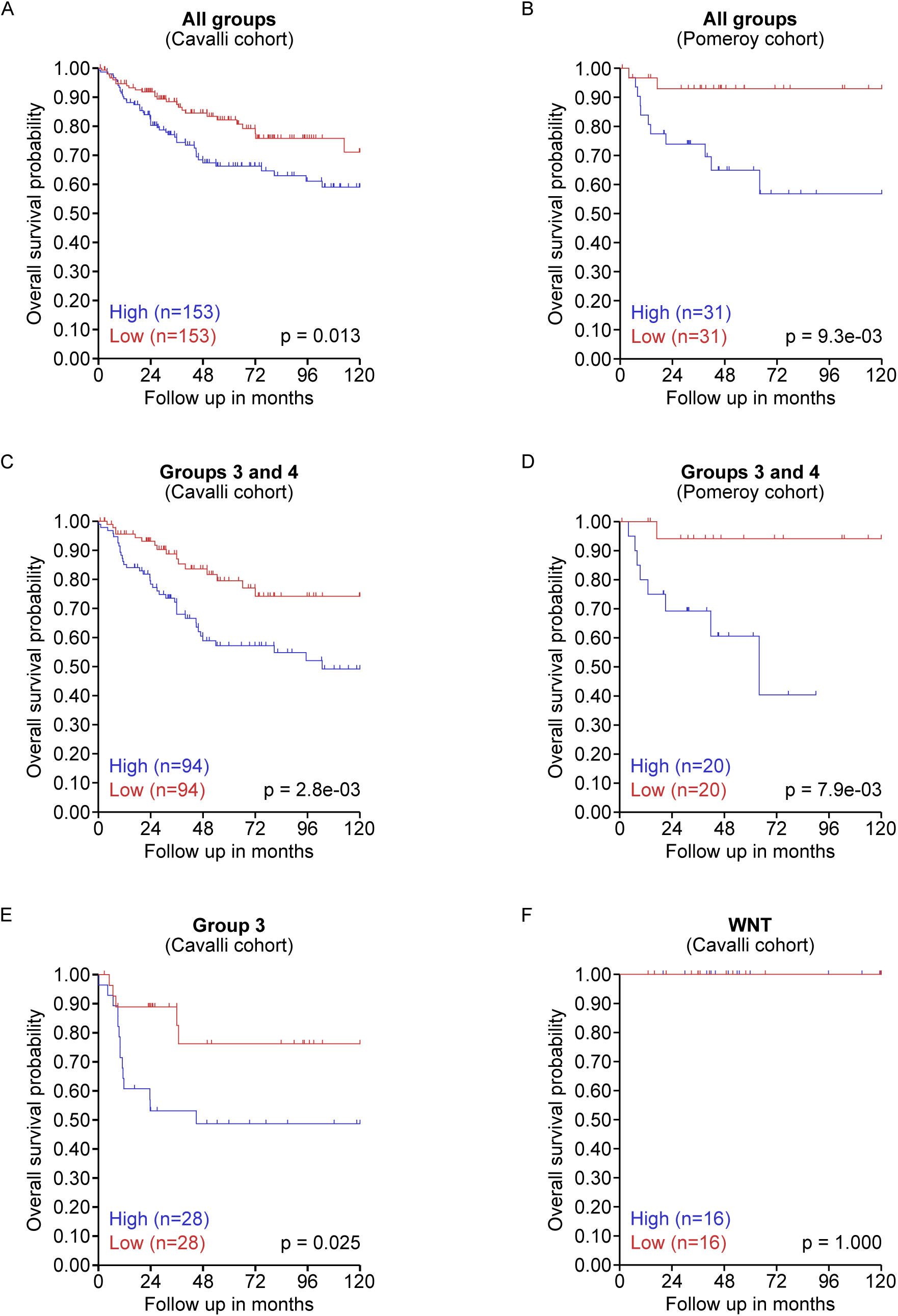
*EIF4EBP1* mRNA expression correlates with overall survival in all MB patients and in Group 3/Group 4 MB patients. **A-F**, Kaplan-Meier survival estimates of overall survival of MB patients stratified by their *EIF4EBP1* mRNA expression levels across all MB patients (A and B), in Group 3 and 4 MB patients combined (C and D), in Group 3 MB patients (E) or in WNT MB patients (F) using data sets from the Cavalli *et al.* [7] and Pomeroy [12] cohorts as indicated. The data were obtained from R2 Genomics and visualization platform and the first versus last quartile was used as cut-off. Significance was calculated with the log-rank test.

### High 4EBP1 protein expression is associated with unfavorable prognosis of MB patients

Since mRNA expression is not strongly correlated with protein expression in MB (Spearman correlation coefficient of 0.53 [20]), we interrogated the prognostic value of 4EBP1 protein expression in this tumor entity. Using a previously established anti-4EBP1 antibody [64], we immunostained FFPE tissue sections from an institutional MB cohort consisting of 61 tumors from all groups, as described previously [17, 62] (Fig. 3A and B). Immunostaining for 4EBP1 was both cytoplasmic and nuclear, consistent with previous reports [22, 48] (Fig. 3B). In line with our observations on *EIF4EBP1* mRNA expression (Supplementary Fig. 1B), we confirmed that 4EBP1 protein levels were higher in relapsed compared to primary MB tissues (Supplementary Fig. 4A). Kaplan Meier analysis showed that positive 4EBP1 staining was strongly associated with reduced overall and progression-free survival across the entire MB cohort (p value < 0.0001; Fig. 3C; p value < 0.0001; Fig. 3D). Additionally, we uncovered that 4EBP1 staining was positively associated with poor overall and progression-free survival in the subset of patients with Group 3 and Group 4 MBs (p value < 0.0001; Fig. 3E; p value = 0.00023; Fig. 3F). Due to the limited number of cases, such correlation could not be determined separately for Group 3 MB patients alone.

**Figure 3.**
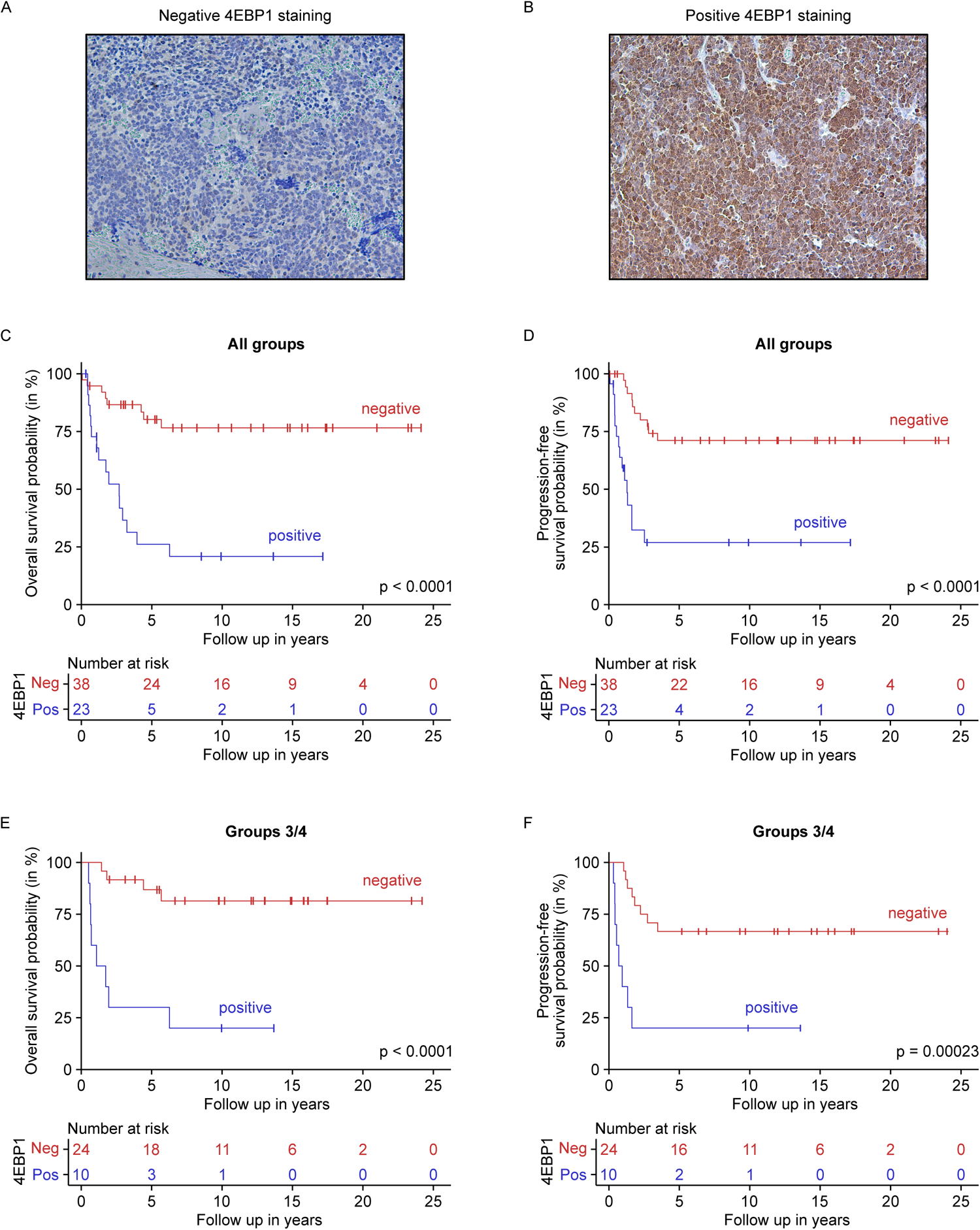
High 4EBP1 protein expression is associated with unfavorable prognosis of MB patients. **A-B**, Representative images of negative (A) and positive (B) 4EBP1 immunohistochemical stainings of selected MB samples represented on the MB TMAs. **C-F**, Kaplan-Meier survival estimates of overall survival (C and E) or progression free-survival (D and F) of MB patients stratified by their 4EBP1 staining score in all patients (C and D) or in Group 3 and Group 4 combined (E and F).

### *EIF4EBP1* expression is regulated at the transcriptional level by MYC in MB

As we uncovered an association between *EIF4EBP1* and *MYC* mRNA expression in MB tissues, and since MYC has been described to control *EIF4EBP1* transcription in other tumor types [2, 60], we asked whether *EIF4EBP1* also represents a MYC target gene in MB. This could provide a molecular mechanism for the *EIF4EBP1* overexpression we observed in the most aggressive MB groups (see Figure 1).

We initially analysed available chromatin immunoprecipitation (ChIP)-sequencing (seq) data from the Encode Consortium, which demonstrated direct binding of MYC at three positions within the *EIF4EBP1* transcriptional regulatory region (encompassing the promoter region, exon 1 and intron 1 [24]) (Fig. 4A). This was detected in various normal and cancer cells, however, not including MB or any type of brain cancer cells. In accordance with previous studies [2, 60], we confirmed the presence of three E boxes, i.e. MYC-binding sites, within this region, including two consensus motifs (CACGTG) and one non-canonical motif (CACATG) (Fig. 4A). Using a luciferase reporter assays covering the nucleotides −192 to +1372 in the *EIF4EBP1* promoter region, exon 1, and part of intron 1 (Fig. 4A), which contains the three ChIP peaks for MYC, we demonstrated that MYC overexpression dose-dependently activated the *EIF4EBP1* promoter in HEK293-T cells (Fig. 4B). To delineate which of the three E boxes are necessary for the transcriptional regulation of the *EIF4EBP1* promoter by MYC, we mutated separately each of the E boxes located within the −192 to +1372 *EIF4EBP1* reporter, as indicated in Fig. 4A. Mutation of only E box 1 compromised MYC-mediated activation of *EIF4EBP1* promoter (Fig. 4C), in contrast to the involvement of the three E boxes for MYCN regulation of *EIF4EBP1* promoter activity as previously reported [64]. These data support that MYC regulates *EIF4EBP1* promoter activity primarily through one specific E box (E box 1), even though it binds three E boxes within this transcriptional regulatory region.

**Figure 4.**
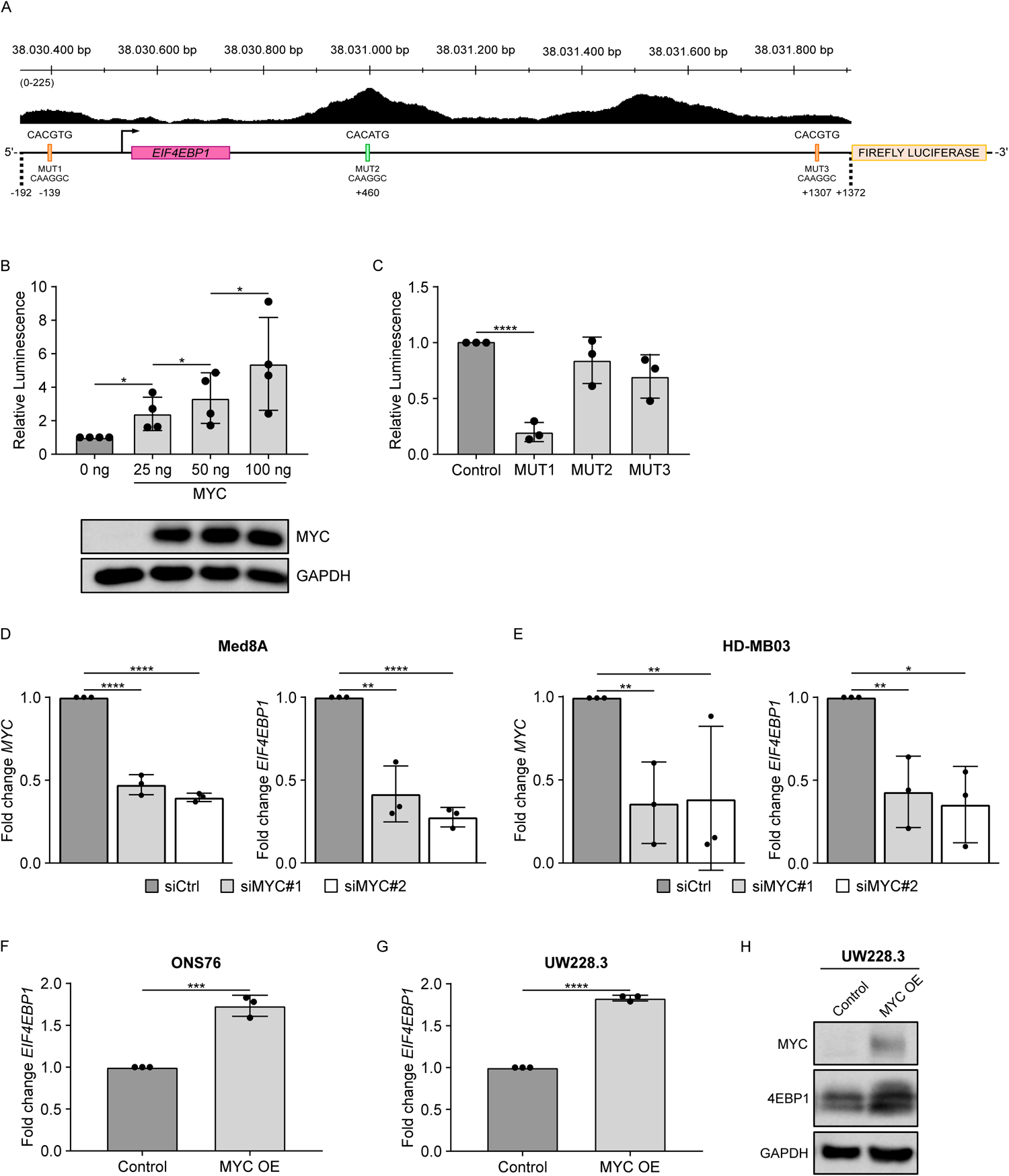
MYC activates *EIF4EBP1* promoter activity and transcription in MB. **A**, ChIP peak locations within the human *EIF4EBP1* promoter, exon 1 and part of intron 1 (hg38; Chr8: 38,030,342 - 38,031,906) from ChIP-sequencing data for MYC (Encode consortium, Encyclopedia of DNA Elements at UCSC [13, 15]); and an illustration of the luciferase reporter construct containing the *EIF4EBP1* promoter, exon 1 and part of intron 1 (−192; +1372), coupled to Firefly luciferase, with the indicated binding sites of the transcription factor MYC. The three E boxes present in the promoter, and corresponding introduced mutations, are indicated. **B and C**, HEK293-T cells were transfected with the (−192; +1372) *EIF4EBP1* promoter reporter construct, together with 25 ng, 50 ng and 100 ng MYC (B) or the (−192; +1372) *EIF4EBP1* promoter reporter constructs containing a mutation of each of the E boxes (as indicated in A), together with 25 ng MYC (C). For B and C, a *Renilla* Luciferase vector was used as an internal control and luciferase activities were detected using the Dual-Luciferase Reporter Assay. Firefly luciferase activity was normalized to Renilla luciferase activity and the ratio was normalized to the corresponding 0 ng (B) or control (C) condition. Data represent the mean of three independent replicates ± standard deviation (SD). Significance was calculated using an unpaired and two-tailed parametric t-test (*p<0.05, ****p<0.0001). A representative immunoblot analyzing expression of MYC is presented in B. **D and E**, Med8A (D) and HD-MB03 (E) MB cells were transiently transfected with negative control siRNAs (siCtrl), or two different siRNAs each targeting MYC (siMYC#1 and siMYC#2). Cells were re-transfected after 96 h with their corresponding siRNA and incubated for a total of 192 h. MRNA was harvested to determine the expression levels of *EIF4EBP1* and *MYC* by qRT-PCR. Data obtained by qRT-PCR represent the mean of three independent replicates ± SD and the fold change in expression was normalized to the negative control (siCtrl). Significance was calculated using an unpaired and two-tailed parametric t-test (*p < 0.05, **p < 0.01, ****p < 0.0001). **F and G**, Control and MYC overexpressing (MYC OE) ONS76 (F) or UW228.3 (G) cells were lysed and levels of *EIF4EBP1* mRNA were determined by qRT-PCR. Data obtained by qRT-PCR represent the mean of three independent replicates ± SD and the fold change in expression was normalized to the control. Significance was calculated using an unpaired and two-tailed parametric t-test (***p<0.001, ****p<0.0001). **H**, Control and MYC overexpressing (MYC OE) UW228.3 cells were lysed and MYC and 4EBP1 levels were determined by immunoblots using the indicated antibodies.

To determine whether MYC is regulating *EIF4EBP1* mRNA expression in MB, we transiently knocked down (KD) MYC in two different *MYC*-amplified MB cell lines, namely Med8A and HD-MB03, and measured *EIF4EBP1* mRNA expression. Upon MYC KD, both *MYC* and *EIF4EBP1* mRNA levels decreased significantly in both cell lines (Fig. 4D and E). To further validate the regulation of *EIF4EBP1*/4EBP1 expression by MYC in MB cells, we used stable MYC overexpression models established in two MB cell lines harbouring low MYC levels, namely ONS76 and UW228.3 [54]. We observed increased *EIF4EBP1* mRNA levels in both MYC overexpressing ONS76 and UW228.3 cells, compared to the corresponding control cells (Fig. 4F and G). Moreover, MYC overexpression resulted in increased 4EBP1 protein levels in UW228.3 cells (Fig. 4H). Taken together, these data support that MYC regulates *EIF4EBP1* transcription in MB by directly regulating its promoter activity.

### 4EBP1 contributes to tumorigenic potential of MB cells

Given the clinical relevance we uncovered for *EIF4EBP1* mRNA and 4EBP1 protein expression in MB patients, we wondered whether 4EBP1 exerts a pro-tumorigenic function in this tumor entity, as reported in gliomas [36] and Ewing sarcomas [21]. Furthermore, we previously reported that 4EBP1 promotes the survival of MB cells under glucose starvation [36], a feature that has been linked to tumorigenic promotion [27]. To investigate the potential tumor-supportive function of 4EBP1 in MB cell models, we investigated HD-MB03 and Med8A cells upon inducible 4EBP1 KD using soft agar colony formation assays. KD of 4EBP1 using two different shRNAs, as validated by 4EBP1 immunoblots, resulted in decreased colony formation of approximately 23% in both MB cell lines when compared to the corresponding shRNA controls (Fig. 5A and B). This indicates that 4EBP1 may contribute to the tumorigenic potential of *MYC*-amplified MB cells.

**Figure 5.**
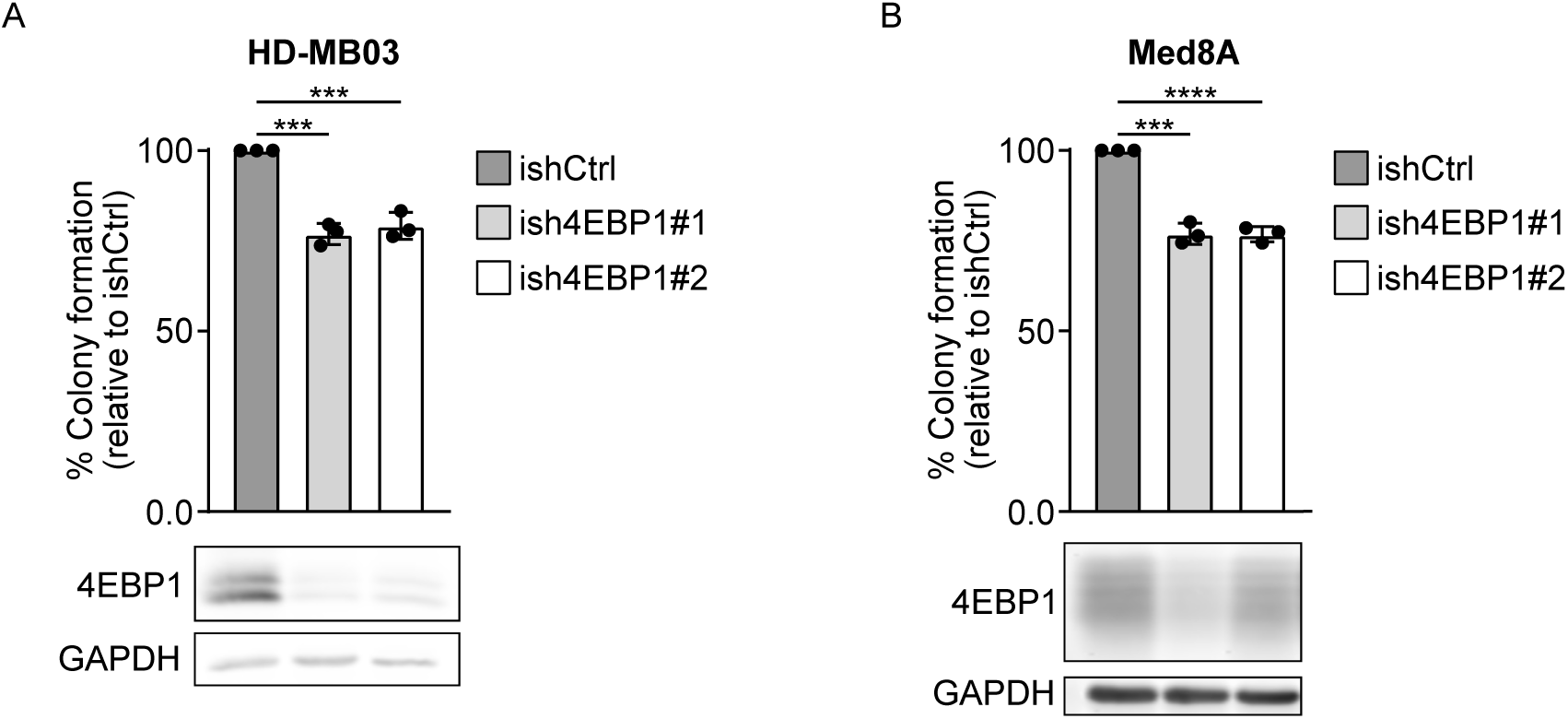
4EBP1 contributes to the tumorigenic potential of *MYC*-amplified MB cell lines. **A and B**, Control (ishCtrl) or stable inducible 4EBP1 knockdown (ish4EBP1#1 and #2) HD-MB03 or (A) Med8A (B) cells were treated with 1 μg/ml doxycycline and grown in soft agar for 21 days. Colonies and single cells were counted, and colony formation efficiency was calculated and normalized to control. Data are reported as means ± SD with indicated significance. Protein expression of 4EBP1 was analyzed by immunoblotting. Significance was calculated using an unpaired and two-tailed parametric t-test (***p<0.001, ****p<0.0001).

## DISCUSSION

We report here that mRNA expression of the mTORC1 substrate and mRNA translational repressor *EIF4EBP1* is increased in MB, as compared to non-neoblastic brain tissue, is higher in Group 3 versus Group 4 MBs, and is associated with patient outcome. This is reminiscent of another negative regulator of mRNA translation, namely *EEF2K*, whose expression in MB was reported to demonstrate the same features [35], suggesting a high level of translational regulation in Group 3 MBs. In line, proteomics analysis revealed that a number of mRNA translation initiation factors, as well as 4EBP1, are overexpressed in Group 3 versus Group 4 MBs [20]. The activity of 4EBP1 is not only dependent on its protein level but also on its phosphorylation states, which is a result of mTORC1 activity [37, 63]. Noteworthy, it was reported that levels of phospho-4EBP1, indicative of inactive 4EBP1, are higher in WNT and SHH as compared to non WNT/non SHH MBs [66]. Additionally, levels of phospho-4EBP1 were similar in Group 3 and Group 4 MBs, while expression of total 4EBP1 protein was higher in Group 3 versus Group 4 MBs [20]. These observations support that 4EBP1 activity is higher in Group 3 MBs compared to other MB groups, likely as a consequence of reduced mTORC1 activity in this MB group. Together with our finding that 4EBP1 protein levels correlate with poor outcome across all MB groups as well as in Group 3/Group 4 MB patients, the available data highlight that *EIF4EBP1* mRNA and 4EBP1 protein expression may represent novel prognostic factors and possible biomarkers in MB.

Amplifications of *MYC* or *MYCN* are well known genetic alterations associated with higher risk MBs [44]. We report that *EIF4EBP1* mRNA expression is associated with *MYC* mRNA expression in MB, an association we found to be particularly evident in Group 3 MBs that are characterized by *MYC* gene amplification [44]. In line with that, we observed that *EIF4EBP1* mRNA expression in Group 4 MBs is associated with elevated *MYCN* mRNA expression, as we previously reported in neuroblastoma [64]. Intriguingly, there was no association between *EIF4EBP1* and *MYC* mRNA levels in WNT MBs. Furthermore, we found fewer MYC target genes to be co-expressed with *EIF4EBP1* in WNT MBs compared to Group 3 MBs, which could reflect differences in MYC activity between the two groups. Interestingly, RNA profiling and proteomics analysis revealed that MYC target genes are highly overexpressed in Group 3 but not in WNT [20], despite similar levels of *MYC* expression in these groups, pointing to a differential activity of MYC in Group 3 versus WNT MBs. One may speculate that transcription factors other than MYC may contribute to *EIF4EBP1* upregulation in WNT MBs, the identity of which remains to be unveiled. Noteworthy, phosphoproteomics data highlighted that MYC phosphorylation as indicator of active MYC is higher in Group 3a MBs ‒ corresponding to subtype II [44] ‒ versus Group 3b and Group 4 MBs [1]. Remarkably, we also uncovered that 4EBP1 protein is more highly expressed in Group 3a MBs compared to Group 3b and Group 4 MBs, further pointing to a link between MYC activity and *EIF4EBP1* mRNA and 4EBP1 protein expression in MB.

Our data indicate that the basis for the *MYC* and *EIF4EBP1* co-expression relies on the transcriptional regulation of *EIF4EBP1* by MYC in MB cells. While it was characterized by ChIP that *EIF4EBP1* is a MYC target gene [2, 60], albeit not in MB cells, it has been elusive whether MYC regulates the *EIF4EBP1* promoter. Here, we provide evidence that MYC activates the *EIF4EBP1* promoter via one E box among the three E boxes previously characterized to be bound by MYC [60]. Additionally, we demonstrated that MYC induces *EIF4EBP1* transcription in MB cell lines as knock down or overexpression of MYC decreased or increased *EIF4EBP1* mRNA levels, respectively. This expands previous studies reporting on the control of *EIF4EBP1* transcription by MYC in colorectal and prostate cancer cells [60], [2].

The function of 4EBP1 in cancer is still under debate as it can exert tumor suppressive or pro-tumorigenic functions [43], depending of the tumor entity and of the metabolic conditions of the tumor microenvironment. For instance, 4EBP1 was shown to mediate cell survival in response to hypoxia and induce angiogenesis in breast cancer models [6]. Furthermore, our previous findings highlighted that 4EBP1 promotes survival of cancer cells, including MB cells, under glucose starvation [36]. As glucose levels are particularly low in MB [3], as compared to other pediatric brain cancers, high 4EBP1 expression may confer resistance to MB cells against such metabolic stress conditions. Since molecular mechanisms of tumor adaptation to glucose starvation are similar to the ones promoting tumorigenesis [27, 36], and since we and others reported that 4EBP1 promotes tumorigenesis of glioblastoma and Ewing’s sarcoma cells *in vivo* [21, 36], we explored 4EBP1 function in MB cell growth *in vitro*. Our findings suggest that 4EBP1 contributes to the tumorigenic capacity, i.e. clonogenic growth in soft agar, of MB cells *in vitro*, albeit to a moderate extent as 4EBP1 knock-down only restrained the clonogenic potential of the investigated MB cells by 20-25%. This could potentially be due to the lack of p53 activity in Med8a and HD-MB03 MB cells, which was reported for Med8a [42], as the pro-tumorigenic properties of 4EBP1 rely on the presence of an intact, active p53 as reported in oncogenic RAS transformed fibroblast models [53]. Noteworthy, the direct protein target of 4EBP1, eIF4E, is a well-known oncoprotein that was shown to promote MB tumorigenesis [32]. Genetic inhibition of eIF4E suppressed MYCN-driven MB development in a genetically engineered mouse model of MB [32]. While this may seem in apparent contradiction with the proposed function of 4EBP1 in MB, it likely reflects the importance of the metabolic conditions within the MB tumor microenvironment. As reported in mouse models of pancreatic cancer and glioblastoma, in well-perfused tumor areas, mTOR and eIF4E are active and their inhibition restricts tumor growth [31, 51]. In contrast, in poorly vascularized tumor areas, mTOR and eIF4E are inactive, while 4EBP1 is activated, which thus facilitates tumor cell survival and favors tumor growth in the long term [31, 51].

Taken together, our findings revealed that elevated *EIF4EBP1* mRNA and 4EBP1 protein expression are associated with shorter survival of MB patients. Increased *EIF4EBP1* mRNA and 4EBP1 protein expression is driven by MYC through direct binding to the *EIF4EBP1* promoter and activation of *EIF4EBP1* transcription, which in turn may contribute to higher clonogenic growth properties of MB cells *in vitro* and possibly more aggressive MB behavior in patients.

## ACKNOWLEDGEMENTS

G.L. was supported by grants from the Deutsche Forschungsgemeinschaft (grant no. LE 3751/2-1), the German Cancer Aid (grant no. 70115129), the Elterninitiative Düsseldorf e.V. (grant no. 701910003), the Dr. Rolf M. Schwiete Stiftung (grant no. 2020-018) and the Research Commission of the Medical Faculty, Heinrich Heine University Düsseldorf (grant no. 2020-044). LH was funded by the Dr. Rolf M. Schwiete Stiftung (grant no. 2020-018).

## SUPPLEMENTARY FIGURE LEGENDS

**Supplementary figure 1.**
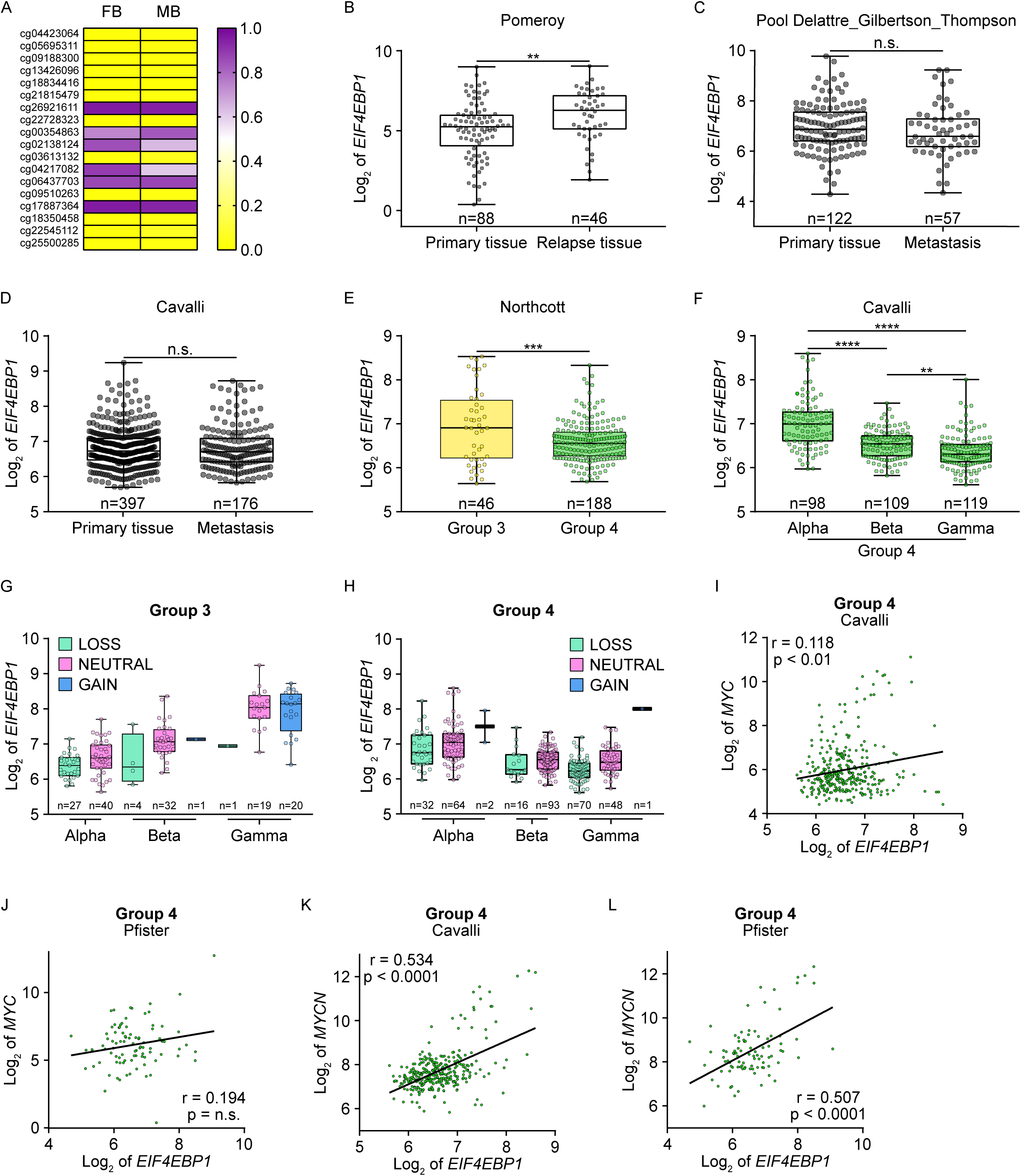
Analyses of *EIF4EBP1* expression levels and *EIF4EBP1* and *MYC(N)* co-expression levels in MB groups and subgroups. **A**, DNA methylation levels of 18 CpG sites located within the *EIF4EBP1* promoter region (human genome GRCh 37/hg19; Chr8: 37,886,955-37,917,868) using the Chatterton dataset for fetal brain (FB) (n=9) [10] and the Cavalli *et al.* dataset for MB tissues (n=763) [7] with 0 representing unmethylated and 1 representing fully methylated CpG sites. A two-tailed Fisher’s exact test was used to determine statistical differences between FB and MB samples. **B**, Expression levels of *EIF4EBP1* mRNA in primary and relapse MB tissues from the Pomeroy [12] cohort. **C and D**, Expression levels of *EIF4EBP1* mRNA in primary and metastatic tissues pooled from the Delattre, Gilbertson [55] and Thompson cohorts (microarray up133p2) (C) and from the Cavalli *et al.* cohort (microarray hugene11t) [7] (D). **E and F**, Expression levels of *EIF4EBP1* levels in Group 3 and Group 4 MBs of the Northcott *et al.* cohort [47] (E) or according to Group 4 MB subgroups of the Cavalli *et al.* cohort [7] (F). Significance in B-F was calculated using an unpaired and two-tailed parametric t-test (**p<0.01, ***p<0.001, ****p<0.0001). **G and H**, Expression levels of *EIF4EBP1* mRNA according to *EIF4EBP1* copy number variation in Group 3 (G) and Group 4 (H) MB subgroups alpha, beta and gamma from the Cavalli *et al.* cohort [7] categorized as *EIF4EBP1* copy number loss (hemizygous deletion [loss]), *EIF4EBP1* neutral copy number (neutral), or *EIF4EBP1* low-level copy number gain (gain). **I-L**, Expression levels of *EIF4EBP1* mRNA in MB patient samples plotted against the mRNA expression levels of *MYC* (I and J) or *MYCN* (K and L) in Group 4 MBs using the Cavalli *et al.* [7] and Pfister [44] cohorts as indicated. Co-expression levels were quantified by calculating the Pearson correlation coefficient.

**Supplementary figure 2.**
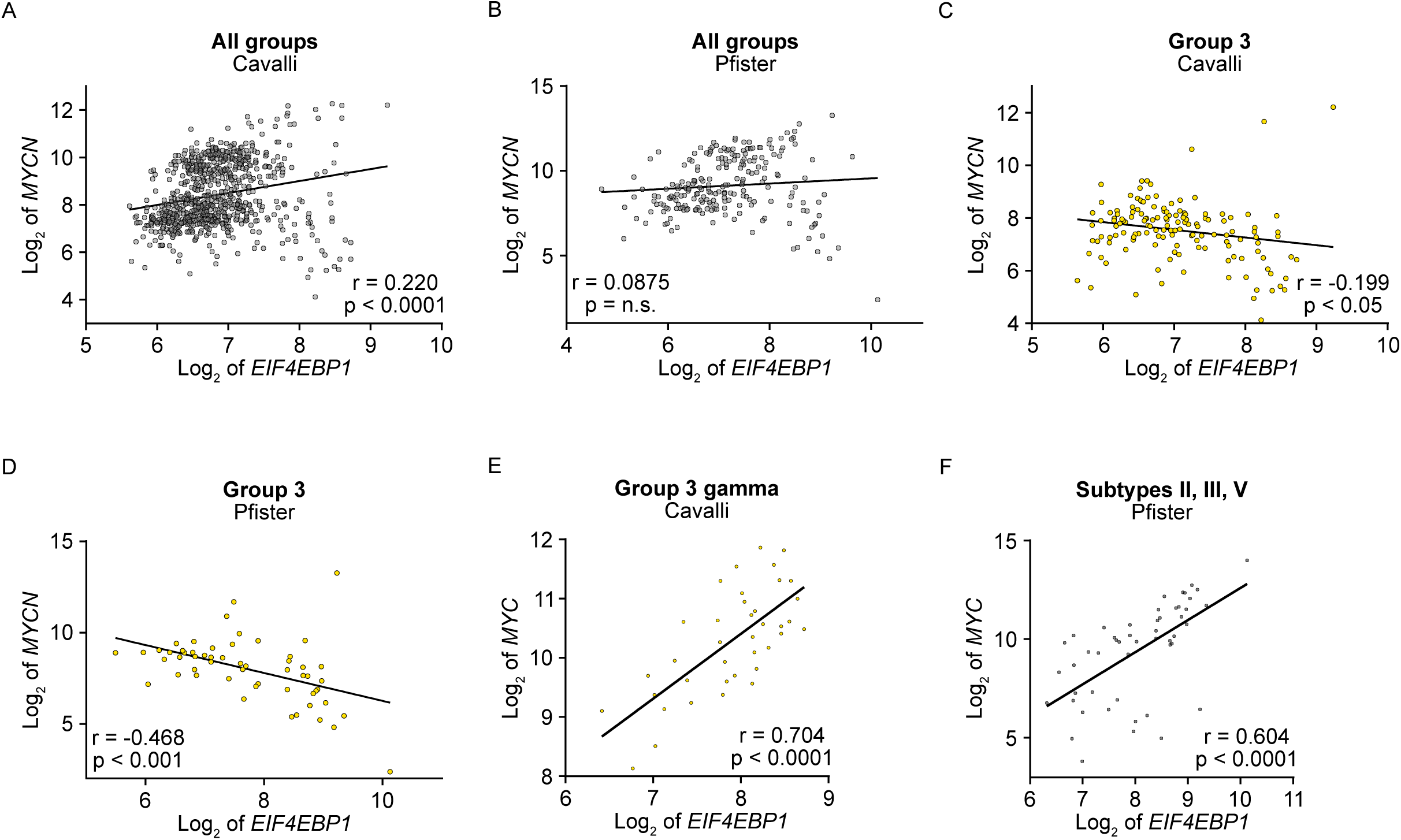
Analyses of *EIF4EBP1* and *MYC(N)* co-expression levels in MB groups and subgroups. **A-D,** Expression levels of *EIF4EBP1* mRNA in MB patient samples plotted against the mRNA expression levels of *MYCN* in all patients (A and B) and in Group 3 MB patients (C and D) using the Cavalli *et al.* [7] and Pfister [44] cohorts as indicated. Co-expression levels were quantified by calculating the Pearson correlation coefficient. **E and F,** Expression levels of *EIF4EBP1* mRNA in MB patient samples plotted against the mRNA expression levels of *MYC* in Group 3 MB gamma (E) and Heidelberg MB subtypes II, III and V (F) using the Cavalli *et al.* [7] and the Pfister [44] cohorts as indicated (see Table S2 for the number of patient samples per group). Co-expression levels were quantified by calculating the Pearson correlation coefficient.

**Supplementary figure 3.**
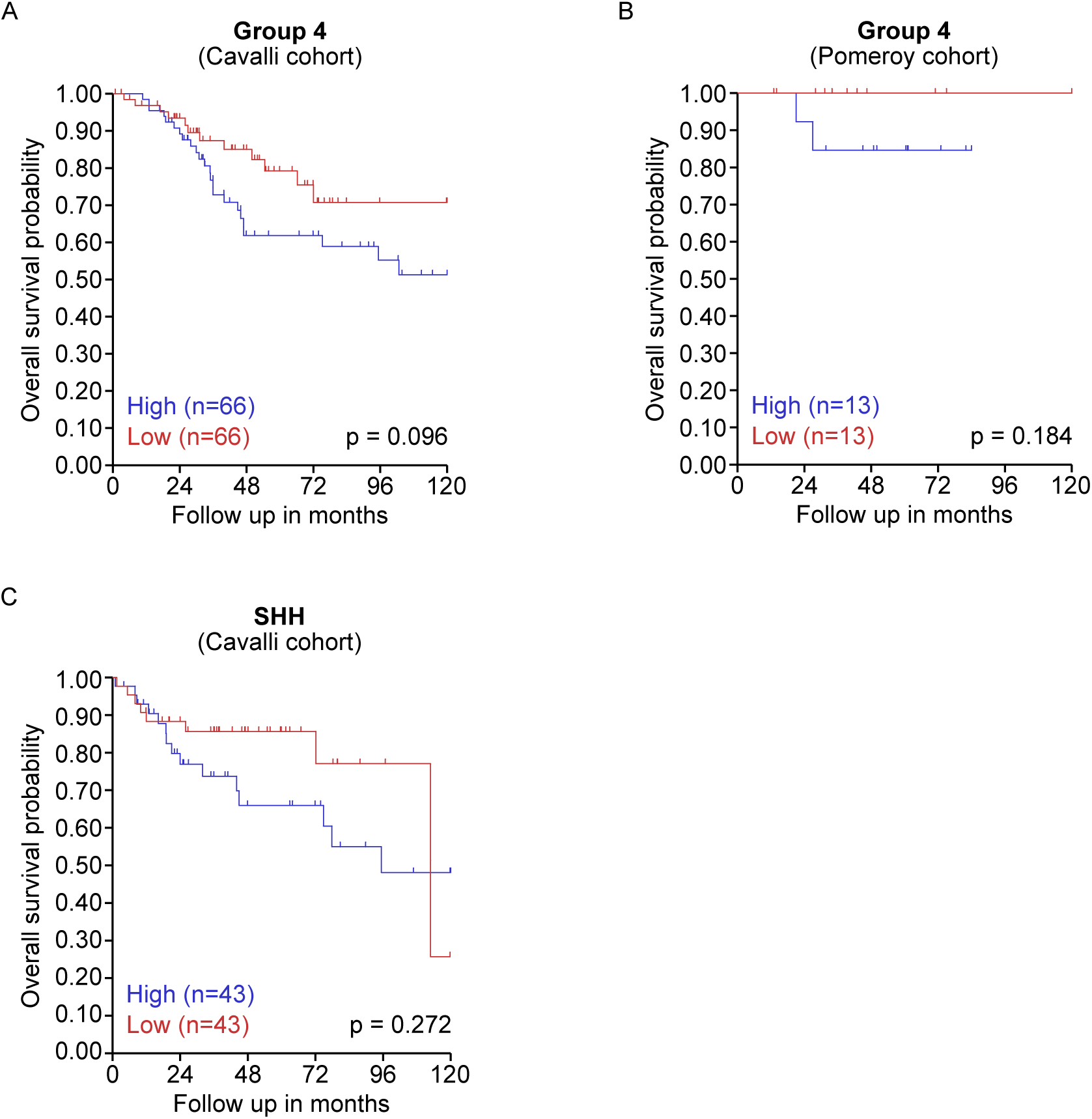
*EIF4EBP1* mRNA expression levels do not correlate with overall survival in Group 4 MB and SHH MB patients. **A-C**, Kaplan-Meier survival estimates of overall survival of MB patients stratified by their *EIF4EBP1* mRNA expression levels in Group 4 (A and B) and in SHH (C) using the Cavalli *et al.* [7] and Pomeroy [12] cohorts as indicated. The data were obtained from R2 Genomics and visualization platform and the median expression level of *EIF4EBP1* mRNA was used as cut-off. Significance was calculated with the log-rank test.

**Supplementary figure 4.**
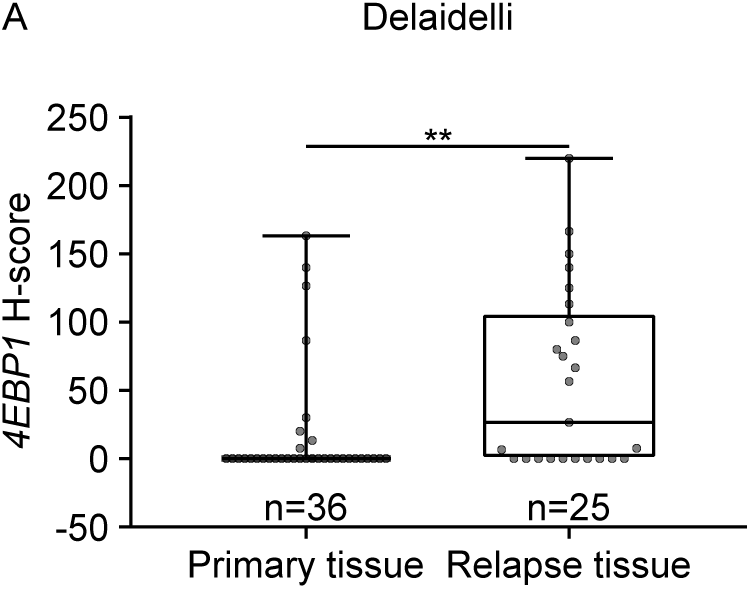
4EBP1 protein expression is upregulated in MB tissue samples from relapsed compared to primary tumors. **A**, Primary and relapsed MB tissue samples were immunostained using an anti-4EP1 antibody. Staining intensity was plotted and significance was calculated using an unpaired and two-tailed parametric t-test (**p < 0.01).

