## Supplementary Tables for "The *EIF4EBP1* gene encoding 4EBP1 is transcriptionally upregulated by MYC and linked to shorter survival in medulloblastoma"

**Supplementary table 1: Overview of the analyzed non-neoplastic brain tissue and medulloblastoma cohorts.**

| Tissue / MB cohorts | Cohort name | Microarray | GEO ID / PUB med link | References |
|---|---|---|---|---|
| **Non-neoplastic brain tissue** | Normal Brain regions - "Berchtold *et al.*" | u133p2 | GSE11882 | [1] |
| | Normal Brain PFC - "Harris *et al.*" | u133p2 | GSE13564 | [5] |
| **Medulloblastoma** | Tumor Medulloblastoma public - "Delattre | u133p2 | Information not available | |
| | Tumor Medulloblastoma Ependymoma - "denBoer“ | u133p2 | GSE74195 | [4] |
| | Tumor Medulloblastoma - "Gilbertson“ | u133p2 | GSE37418 | [10] |
| | Tumor Medulloblastoma - ATRT - "Hsieh“ | u133p2 | GSE67851 | [6] |
| | Tumor Medulloblastoma PLoS One - "Kool *et al.*" | u133p2 | GSE10327 | [7] |
| | Tumor Medulloblastoma - "Pfister“ | u133p2 | 28726821 | [8] |
| | Mixed Medulloblastoma public - "Pomeroy” | u133a | 21098324 | [3] |
| | Tumor Medulloblastoma - "Thompson” | u133a | Information not available | |
| | Tumor Medulloblastoma - "Cavalli *et al.*" | hugene11t | GSE85217 | [2] |
| | Tumor Medulloblastoma MAGIC - "Northcott *et al.*" | hugene11t | GSE37382 | [9] |

**Supplementary table 2: Overview of the number of patients per subgroup in the different cohorts.**

| MB cohorts | MB group | Number of patients |
|---|---|---|
| **"Cavalli *et al.*"** | **All MB groups** | **763** |
| | SHH | 223 |
| | WNT | 70 |
| | Group 3 | 144 |
| | Group 4 | 326 |
| **"Gilbertson”** | SHH | 10 |
| | WNT | 8 |
| | Group 3 | 16 |
| | Group 4 | 39 |
| **"Kool *et al.*"** | SHH | 15 |
| | WNT | 9 |
| | Group 3 | 11 |
| | Group 4 | 27 |
| **"Northcott *et al.*"** | Group 3 | 46 |
| | Group 4 | 188 |
| **"Pfister”** | All MB groups | 223 |
| | SHH | 59 |
| | WNT | 17 |

| | | |
|---|---|---|
| | Group 3 | 56 |
| | Group 4 | 91 |
| **"Pomeroy”** | SHH | 52 |
| | WNT | 14 |
| | Group 3 | 51 |
| | Group 4 | 71 |

**Supplementary table 3: Chromosomal positions of the CpG sites within the *EIF4EBP1* promoter region Chr8: 37,886,955-37,917,868 (human genome GRCh 37/hg19).**

| CpG site (ID) | Chromosomal location |
|---|---|
| cg00354863 | Chr8:37886955 |
| cg22545112 | Chr8:37887635 |
| cg22728323 | Chr8:37887713 |
| cg09510263 | Chr8:37887715 |
| cg21815479 | Chr8:37887900 |
| cg09188300 | Chr8:37887903 |
| cg04423064 | Chr8:37887926 |
| cg05695311 | Chr8:37887949 |
| cg18350458 | Chr8:37887952 |
| cg13426096 | Chr8:37887990 |
| cg18834416 | Chr8:37888184 |
| cg25500285 | Chr8:37888493 |
| cg03613132 | Chr8:37888764 |
| cg04217082 | Chr8:37889752 |
| cg17887364 | Chr8:37891957 |
| cg02138124 | Chr8:37901697 |
| cg06437703 | Chr8:37914619 |
| cg26921611 | Chr8:37917868 |

**Supplementary table 4: List of siRNA sequences.**

| Target gene and siRNA name | siRNA sequence |
|---|---|
| Dharmacon | |
| Non-targeting | 5’- UAAGGCUAUGAAGAGAUAC -3’ |
| | 5’- AUGUAUUGGCCUGUAUUAG -3’ |
| | 5’- AUGAACGUGAAUUGCUCAA -3’ |
| | 5’- UGGUUUACAUGUCGACUAA -3’ |
| MYC si14 | 5’- AACGUUAGCUUCACCAACA -3’ |
| MYC si35 | 5’- CUACCAGGCUGCGCGCAAA -3’ |

**Supplementary table 5: List of RT-qPCR primer sequences.**

| Target transcript | Primer sequence |
|---|---|
| *4EBP1* | FW: 5’-AGCCCTTCCAGTGATGAGC-3’<br>RV: 5’-TGTCCATCTCAAACTGTGACTCTT-3’ |
| *GusB* | FW: 5’-GTTTTTGATCCAGACCCAGATG-3’<br>RV: 5’-GCCCATTATTCAGAGCGAGTA-3’ |
| *MYC* | FW: 5’-GTCAAGAAGCGAACACACAAC-3’<br>RV: 5’-TTGGACGGACAGGATGTATGC-3’ |
| *PPIA* | FW: 5’-TTATTTGGGTTGCTCCCTTC-3’<br>RV: 5’-AAGTGTGCCAAATCTGCAAG-3’ |

| | |
|---|---|
| *ACTB* | FW: 5'-TCCCCCAACTTGAGATGTATG-3'<br>RV: 5'-ACTGGTCTCAAGTCAGTGTACAGG-3' |

**Supplementary table 6: List of antibodies used for immunoblots.**

| Antibody | Company | Catalog number |
|---|---|---|
| 4EBP1 (53H11) | Cell signaling, Cambridge, UK | #9644S |
| Anti-rabbit IgG, HRP linked antibody | Cell signaling | #7074 |
| GAPDH (14C10) | Cell signaling | #2118S |
| IRDye® 800CW Goat anti-Mouse IgG Secondary Antibody | LI-COR Bioscience, Bad Homburg, Germany | #925-32210 |
| IRDye® 800CW Goat anti-Rabbit IgG Secondary Antibody | LI-COR Bioscience | #925-32211 |
| MYC | Cell signaling | #5605 |
| β-ACTIN | Sigma Aldrich, St Louis, USA | #A2228 |

**Supplementary table 7: Results of pair-wise comparison of *EIF4EBP1* expression levels between different MB groups in the investigated cohorts (corresponding to Fig. 1B and C).**

| MB Cohort | MB group comparisons | Significance |
|---|---|---|
| **"Cavalli *et al.*"** | Group 3 vs. SHH | ** |
| | Group 3 vs. WNT | n.s. |
| | Group 3 vs. Group 4 | **** |
| | WNT vs. SHH | **** |
| | WNT vs. Group 4 | **** |
| | SHH vs. Group 4 | **** |
| **Cohort pool (Pfister, Gilbertson, Pomeroy, Kool *et al.*)** | Group 3 vs. SHH | n.s. |
| | Group 3 vs. WNT | n.s. |
| | Group 3 vs. Group 4 | **** |
| | WNT vs. SHH | n.s. |
| | WNT vs. Group 4 | **** |
| | SHH vs. Group 4 | **** |
